# IGF-I Governs Cortical Inhibitory Synaptic Plasticity By Astrocyte Activation

**DOI:** 10.1101/2020.02.11.942532

**Authors:** José Antonio Noriega-Prieto, Laura Eva Maglio, Jonathan A. Zegarra-Valdivia, Jaime Pignatelli, Ana M. Fernandez, Laura Martinez-Rachadell, Jansen Fernandes, Ángel Núñez, Alfonso Araque, Ignacio Torres Alemán, David Fernández de Sevilla

## Abstract

Insulin-like growth factor-I (IGF-I) signaling plays key regulatory roles in multiple processes of brain physiology and pathology. While the direct effects of IGF-I in neurons have been extensively studied, the astrocyte involvement in IGF-I signaling and the consequences on synaptic plasticity and animal behavior remain unknown. Here we show that IGF-I induces the long-term depression (LTD) of inhibitory synaptic transmission in the mouse barrel cortex. This LTD requires the activation of the IGF-I receptor (IGF-IR) in astrocytes, which stimulates astrocyte Ca^2+^ signaling and the release of ATP/adenosine that in turn activates A_2A_ adenosine receptors at presynaptic inhibitory terminals. Specific deletion of IGF-IR in cortical astrocytes (IGF-IR^-/-^) impaired the behavioral performance in a whisker discrimination task. These results show novel mechanisms and functional consequences of IGF-I signaling on cortical inhibitory synaptic plasticity and animal behavior, revealing astrocytes as key elements in these processes.

## INTRODUCTION

Insulin-like growth factor-I (IGF-I) is a peptide with well-known trophic functions. IGF-I is actively transported to the central nervous system (CNS) from plasma through the blood-brain barrier (1)(2), and is also locally produced by neurons and glial cells (3)(4)(5). IGF-I regulates neuronal firing (6)(7) and modulates excitatory synaptic transmission in many areas of the central nervous system (8)(9)(10)(11)(12). IGF-I also produces a long-lasting depression of glutamate-mediated GABA release by Purkinje cells in the cerebellum (10), or a long-term potentiation of GABA release in the olfactory bulb (13). However, whether IGF-I modulates inhibitory synaptic transmission in the neocortex remains unexplored.

Astrocytes have emerged as active elements directly involved in synaptic physiology. They respond with Ca^2+^ elevations to neurotransmitters released by neurons and induce changes in neuronal excitability and synaptic transmission by releasing gliotransmitters (14)(15)(16)(17)(18)(19)(20). Astrocytes can release a variety of trophic factors that promote neuronal survival, including IGF-I (21), and several studies have shown the presence of IGF-I and IGF-IRs in neurons, astrocytes and microglia (22)(23)(24)(5). Indeed, IGF-I regulates astrocytic glucose control and CNS glucose metabolism (25), exerts proliferative effects on astrocytes (26), and reduces their cAMP levels(27). Activation of GABA_B_Rs in astrocytes induce release of glutamate that potentiates both inhibitory (28) and excitatory (29) synaptic transmission. In addition to glutamate, astrocytes release ATP that in turn depresses excitatory synaptic transmission in the hippocampus (30)(31)(32). Moreover, in the neocortex, exocytosis of ATP from astrocytes leads to a short term down regulation of inhibitory synaptic currents by inhibiting postsynaptic and extrasynaptic GABA_A_ receptors in layer 2/3 pyramidal neurons (33).

Here, we investigated whether IGF-IR activation on astrocytes induces long-term modulation of inhibitory synaptic transmission at layer 2/3 pyramidal neurons of the barrel cortex. We found that IGF-I induces a presynaptic LTD of IPSCs that depends on cytosolic calcium increases in astrocytes, and activation of A_2A_ adenosine receptors. This LTD of inhibitory synaptic transmission is absent in mice in which IGF-IR has been deleted specifically in astrocytes (IGF-IR^-/-^ mice). In addition, we demonstrate that these IGF-IR^-/-^ mice show an impairment in the performance of a whisker discrimination task. Therefore, our results demonstrate a novel mechanism of long-term synaptic depression of inhibition at the barrel cortex induced by activation of astrocytic IGF-IRs and ATP/Adenosine (ATP/Ado) release from astrocytes that have important consequences in the processing of somatosensory information occurring during the whisker discrimination task.

## RESULTS

### IGF-I induces presynaptic LTD of IPSCs (iLTD_IGFI_) that requires activation of astrocytes

We first investigated whether IGF-I modulates the efficacy of inhibitory synaptic transmission at layer 2/3 pyramidal neurons of the barrel cortex. Inhibitory postsynaptic currents (IPSCs) evoked by stimulation of layer 4 were recorded in layer 2/3 neurons. IPSCs were isolated in the presence of the glutamate receptor antagonists 20 μM CNQX and 50 μM AP5 (**Figure 1A**). After 5 min of IPSC recording, IGF-I (10 nM) was bath applied during 35 min and then washed-out (**Figure 1B**). IGF-I induced a long-term depression (LTD) of the IPSCs peak amplitude (termed iLTD_IGFI_) that persisted after IGF-I washout (from 100.6 ± 1.42 to 60.73 ± 5.08 % of IPSC peak amplitude, before and after IGF-I. N = 7, P < 0.001; **Figure 1B**, black circles). These IGF-I-induced effects were prevented by bath perfusion of the IGF-IR selective inhibitor NVP-AEW 541 (from 98.72 ± 0.50 to 96.15 ± 3.59 % of IPSC peak amplitude, before and after IGF-I. N = 7, P = 0.50; **Figure 1B**, white circles), indicating that were mediated by IGF-IR activation.

**Figure 1.**
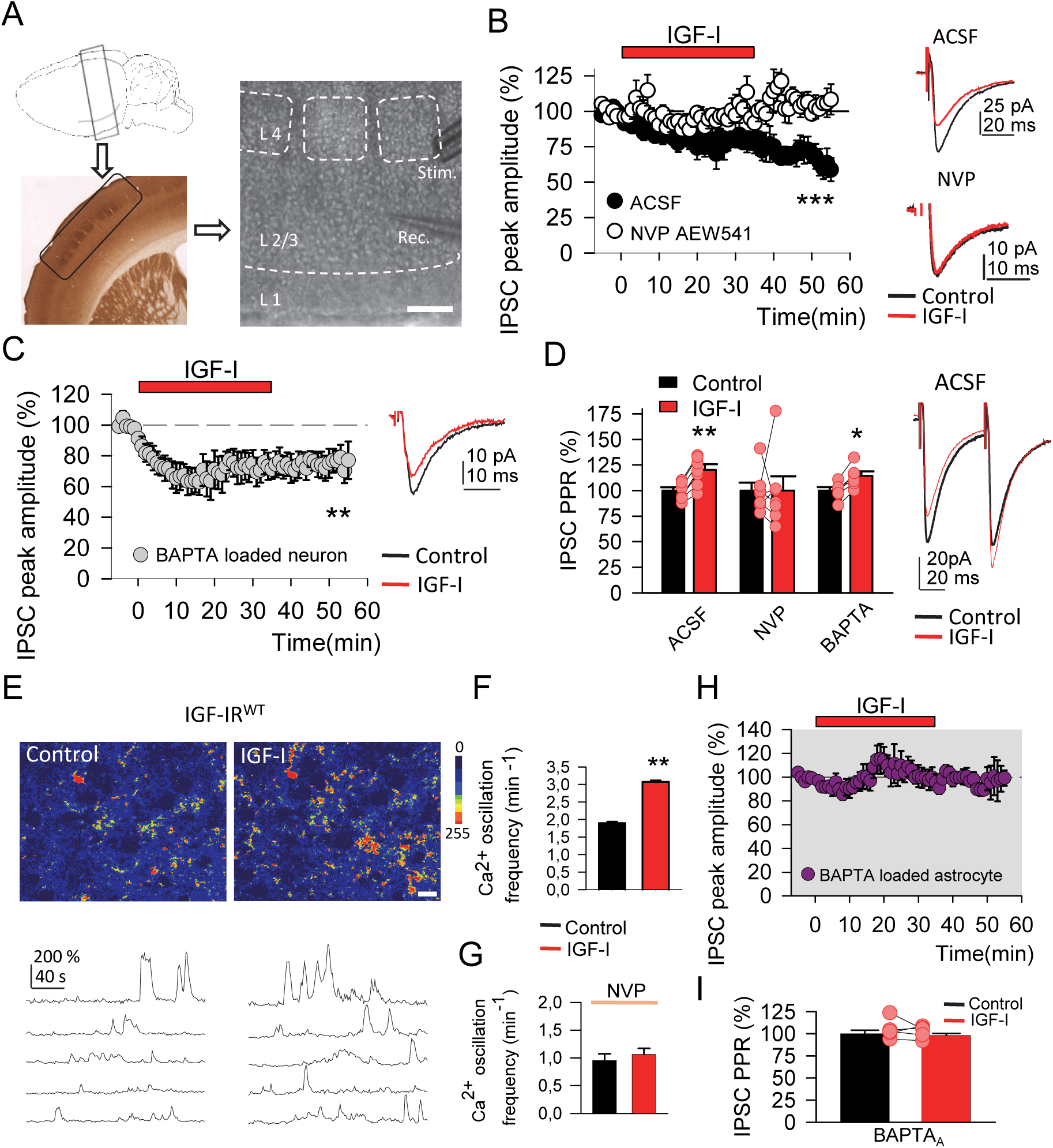
IGF-I mediated Astrocyte signaling induces LTD of IPSCs. ***A. left.*** Representative drawing of location of brain section (top) and slice treated with cytochrome oxidase (bottom) of barrel cortex. ***Right.*** DIC image of slice showing the stimulation (stim) and recording (rec) electrodes (scale bar, 100 µm). ***B. Left.*** Time course of IPSCs peak amplitude in ACSF (black circles) and in presence of NVP (white circles) showing the effect of IGF-I bath application. ***Right.*** Representative IPSCs in ACSF and NVP before and during IGF-I. ***C. Left.*** Same as B left in ACSF but with intracellular BAPTA. ***Right.*** Same as B right in ACSF but with intracellular BAPTA. ***D. Left.*** PPR of the IPSCs in ACSF, NVP and intracellular BAPTA. ***Right.*** Representative IPSCs evoked by paired pulse stimulation in ACSF before and during IGF-I. ***E. Top.*** Pseudocolor Ca^2+^ images showing the intensities of GCaMP6f expressing astrocytes in barrel cortex, before and during IGF-I application in slices from IGF-IR^WT^ mice (scale bar, 50 µm). ***Bottom.*** Representative calcium traces in astrocytes from IGF-IR^WT^ mice. ***F.*** Bar plot of the calcium oscillation frequency in astrocytes from IGF-IR^WT^ mice before and during IGF-I. ***G.*** Same as H but in the presence of NVP. ***H.*** Same as B left in ACSF but after 30 min of BAPTA loaded astrocyte. ***I.*** Same as D left in ACSF but after 30 min of BAPTA loaded astrocyte.

Because postsynaptic calcium increases are known to be required in the induction of long-term synaptic plasticity (34), we then tested whether iLTD_IGFI_ induction required cytosolic calcium elevation in the recorded neuron. We used the same experimental approach described above, but now including the Ca^2+^chelator BAPTA (40 mM) in the patch pipette to prevent neuronal calcium elevations. BAPTA loading of the postsynaptic neuron abolished the calcium increases evoked by neuronal depolarization (**Supplementary Figure 1**). However, a similar iLTD_IGFI_ was induced in both control and neuron BAPTA-loaded conditions (from 100.08 ± 0.60 to 68.67 ± 6.76 % of IPSC peak amplitude, before and after IGF-I. N = 7, P < 0.01; **Figure 1C**). Taken together, these data demonstrate that IGF-I induces LTD of IPSCs without requiring cytosolic calcium elevations in the postsynaptic neuron. To test a possible presynaptic origin of the effect of IGF-I, we recorded the IPSCs evoked by paired-pulse stimulation (50 ms delay), and analyzed changes in paired-pulse responses (PPRs). PPRs were increased during IGF-I-mediated LTD of the IPSCs (from 100.00 ± 3.23 to 120.37 ± 5.33 % of IPSC paired-pulse responses, before and after IGF-I. N = 7, P < 0.01; **Figure 1D**, ACSF) indicating that they were mediated by a presynaptic mechanism.

Astrocytes are emerging as important cells involved in the regulation of synaptic transmission and plasticity(35). Therefore, we next determined whether iLTD_IGFI_ requires cytosolic calcium elevations in astrocytes. First, we tested whether astrocytes responded to IGF-I blocking glutamatergic, GABAergic, cholinergic, purinergic and dopaminergic receptors. We used a cocktail containing CNQX and D-AP5 (20 μM and 50 μM; for AMPA/kainite and NMDA glutamate receptors respectively), MPEP and LY367385 (50 μM and 100 μM; for mGluR5 and mGluR1 respectively), picrotoxin and CGP (50 μM and 1μM for GABA_A_Rs and GABA_B_Rs respectively), atropine (50 μM; for muscarinic cholinergic receptors), CPT (2 μM; for A1 adenosine receptors), suramin (100 μM; for P2 purinergic receptors) and flupenthixol (30 μM; for D1/D2 dopaminergic receptors). In addition, TTX (1 μM) was also added to the cocktail to prevent action potential-mediated neurotransmitter release. Application of IGF-I induced an increase in the frequency of astrocyte calcium elevations (from 0.75 ± 0.08 to 1.22 ± 0.10 min^-1^ before and during IGF-I. N = 46 astrocytes, P < 0.01; **Figure 3C**, ACSF. From 1.60 ± 0.03 to 2.93 ± 0.05 min^-1^ before and during IGF-I. N = 133 processes, P < 0.01; **Figure 1E, F**), that was absent under NVP (from 0.89 ± 0.14 to 0.97 ± 0.14 min^-1^ before and during IGF-I. N = 81 astrocytes, P = 0.73; **Figure 1G**, NVP). Then, we tested whether the IGFI-induced enhancement of astrocyte calcium signal contributed to the iLTD_IGFI_. We prevented the Ca^2+^ signal selectively in astrocytes, by recording a cortical astrocyte with a patch pipette containing 40 mM BAPTA. Since cortical astrocytes are gap-junction connected, BAPTA injected in a single astrocyte can diffuse throughout a large extension of the gap junction-coupled astrocytic network (30)(36)(37). After BAPTA-loading of the astrocytic network, IGF-I failed to modulate the IPSCs (from 99.79 ± 0.62 to 92.75 ± 6.20 % of IPSC peak amplitude, before and after IGF-I. N = 6, P = 0.31; **Figure 1H**) or the PPR (from 100.00 ± 3.86 to 98.01 ± 2.48 % of IPSC paired-pulse responses, before and after IGF-I. N = 6, P = 0.67; **Figure 1I**). Taken together, these data suggest that cytosolic Ca^2+^ increases in astrocytes, but not in the recorded PN (see above results), are essential for the induction of iLTD_IGFI_.

### iLTD_IGFI_ requires IGF-IR activation in astrocytes

Above experiments show that iLTD_IGFI_ requires astrocyte calcium elevations. However, astrocyte stimulation by IGF-I could occur directly through either activation of astrocytic receptors, or indirectly through activation of neuronal receptors that trigger an indirect signaling pathway. To discriminate between these two possibilities, we deleted IGF-IR specifically in astrocytes using a combined viral and genetic approach. Mice with floxed IGF-IR gene (IGF-IR^f/f^ mice) were injected in the barrel cortex with the virus AAV8-GFAP-Cre-mCherry (or AAV8-GFAP-mCherry as a control) that included the Cre-recombinase under the astroglial promoter GFAP, and mCherry as a reporter (**Figure 2A)**. To delete IGF-IRs selectively in cortical astrocytes, mice were injected in the barrel cortex with 500 nl of the viral vector AAV8-GFAP-Cre-mCherry (or AAV8-GFAP-mCherry as a control, i.e., lacking the Cre recombinase). Immunohistochemistry analysis confirmed the selective expression of the virus in cortical astrocytes. Furthermore, in IGF-IR^f/f^ mice injected with AAV8-GFAP-Cre-mCherry, mCherry expression was largely reduced and confined to the soma (**Figure 2B, mCherry, IGF-IR^-/-^**) compared to the higher and more spread signal in control littermates (**Figure 2B, mCherry, IGF-IR^WT^**).

**Figure 2.**
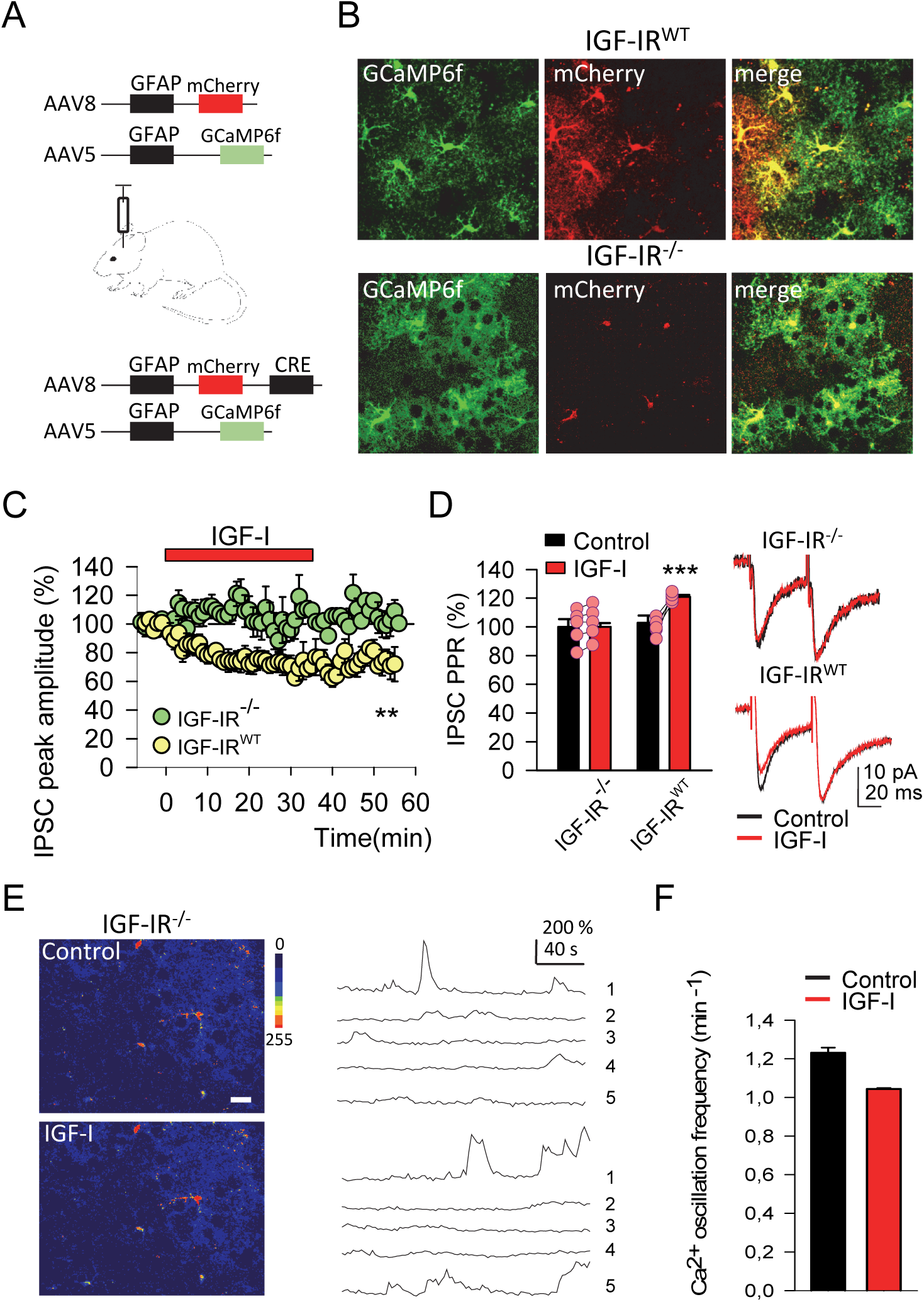
Removal of IGF-IR in astrocytes avoid the iLTD_IGFI_. ***A.*** Schema of experimental procedure showing the local injection of GFAP-mCherry-CRE (or AAV8-GFAP-mCherry as a control) and GFAP-GCaMP6f. ***B.*** Representatives images of locally targeting astrocytes with GFAP-GCaMP6f and GFAP-mCherry for IGF-IR^WT^ (upper images, green and red images), and GFAP-GCaMP6f and GFAP-mCheryy-CRE for IGF-IR^-/-^ (lower images, green and red images). ***C.*** Same as 1B left in ACSF but in IGF-IR^-/-^ and IGF-IR^WT^ mice (green and yellow circles respectively). ***D. Left.*** PPR of the IPSCs in IGF-IR^-/-^ and IGF-IR^WT^ mice before and during IGF-I application. ***Right.*** Representative IPSCs evoked by paired pulse stimulation in IGF-IR^-/-^ and IGF-IR^WT^ mice before and during IGF-I. ***E. Left.*** Same as 1G Top but in IGF-IR^-/-^ mice (scale bar, 50 µm). ***Right.*** Representative calcium traces in astrocytes from IGF-IR^-/-^ mice. ***F.*** Same as 1H but from IGF-IR^-/-^ mice.

As expected, in IGF-IR^WT^, IGF-I elevates astrocyte calcium levels (**Figure 1E, F**) inducing the iLTD_IGFI_ and increasing the PPR (from 99.89 ± 0.80 to 74.74 ± 4.97 % of IPSC peak amplitude before and after IGF-I. N = 5, P < 0.01; **Figure 2C**, yellow circles; PPR: from 100.00 ± 2.64 to 120.92 ± 1.31 % of IPSC paired-pulse responses, before and after IGF-I. N = 5, P < 0.001; **Figure 2D**, IGF-IR^WT^). In contrast, in IGF-IR^-/-^ mice, IGF-I did not modify the frequency of calcium elevations in astrocytes (from 1.23 ± 0.02 to 1.04 ± 0.004 min^-1^ before and during IGF-I. N = 67 processes, P = 0.28; **Figure 2E, F**) and did not alter the IPSC amplitude and PPR (from 100.27 ± 1.55 to 107.16 ± 5.62 % of IPSC peak amplitude before and after IGF-I. N = 5, P = 0.29; **Figure 2C**, green circles; PPR: from 100.00 ± 5.36 to 102.92 ± 5.05 % of IPSC paired-pulse responses, before and after IGF-I. N = 5, P = 0.70; **Figure 2D**, IGF-IR^-/-^). These results demonstrate that IGF-IR in astrocytes are necessary for the induction of iLTD_IGFI_.

### iLTD_IGFI_ requires A_2A_ adenosine receptor activation

Astrocytic activation may stimulate the release of ATP, which, after being converted to adenosine, may regulate synaptic transmission(38), (35) in the hippocampus, cortex and striatum(39), (40), (41). Therefore, we tested whether astrocytic ATP/Adenosine was responsible for the iLTD_IGFI_. The iLTD_IGFI_ was abolished by the antagonist of adenosine A_2A_ receptors SCH 58261 (100 nM; from 100.02 ± 0.63 to 94.98 ± 2.43 % of IPSC peak amplitude, before and after IGF-I. N = 8, P = 0.07; **Figure 3A**, purple circles), but not by the antagonist of adenosine A1 receptors, CPT (5 µM; from 98.72 ± 1.75 to 55.74 ± 7.08 % of IPSC peak amplitude, before and after IGF-I. N = 7; P < 0.001; **Figure 3A**, blue circles). Moreover, the increase in PPR induced by IGF-I was preserved in the presence of CPT but absent in SCH 58261 (from 100.00 ± 5.90 to 118.42 ± 4.95 % of IPSC paired-pulse responses in the presence of CPT, before and after IGF-I. N = 7, P < 0.05. From 100.00 ± 2.96 to 99.76 ± 1.85 % of IPSC paired-pulse responses in the presence of SCH, before and after IGF-I. N = 8, P = 0.94; **Figure 3B**). To test whether adenosine-receptor activation occurs downstream of astrocytic calcium activity, we analyzed the effects of A_2A_ and A_1_ receptor antagonists on astrocytic calcium signals. We observed that IGF-I evoked an increase in calcium event frequency in the presence of SCH (from 0.67 ± 0.12 to 1.40 ± 0.17 min^-1^ before and during IGF-I. N = 48 astrocytes, P < 0.01; **Figure 3C**, SCH), and CPT (from 0.59 ± 0.10 to 1.20 ± 0.13 min-1 before and during IGF-I. N = 72 astrocytes, P < 0.01; **Figure 3C**, CPT). Taken together, these results indicate that IGF-I induces an increase in calcium levels in astrocytes, which leads to the activation of A_2A_ receptors, thus inducing iLTD_IGFI_.

**Figure 3.**
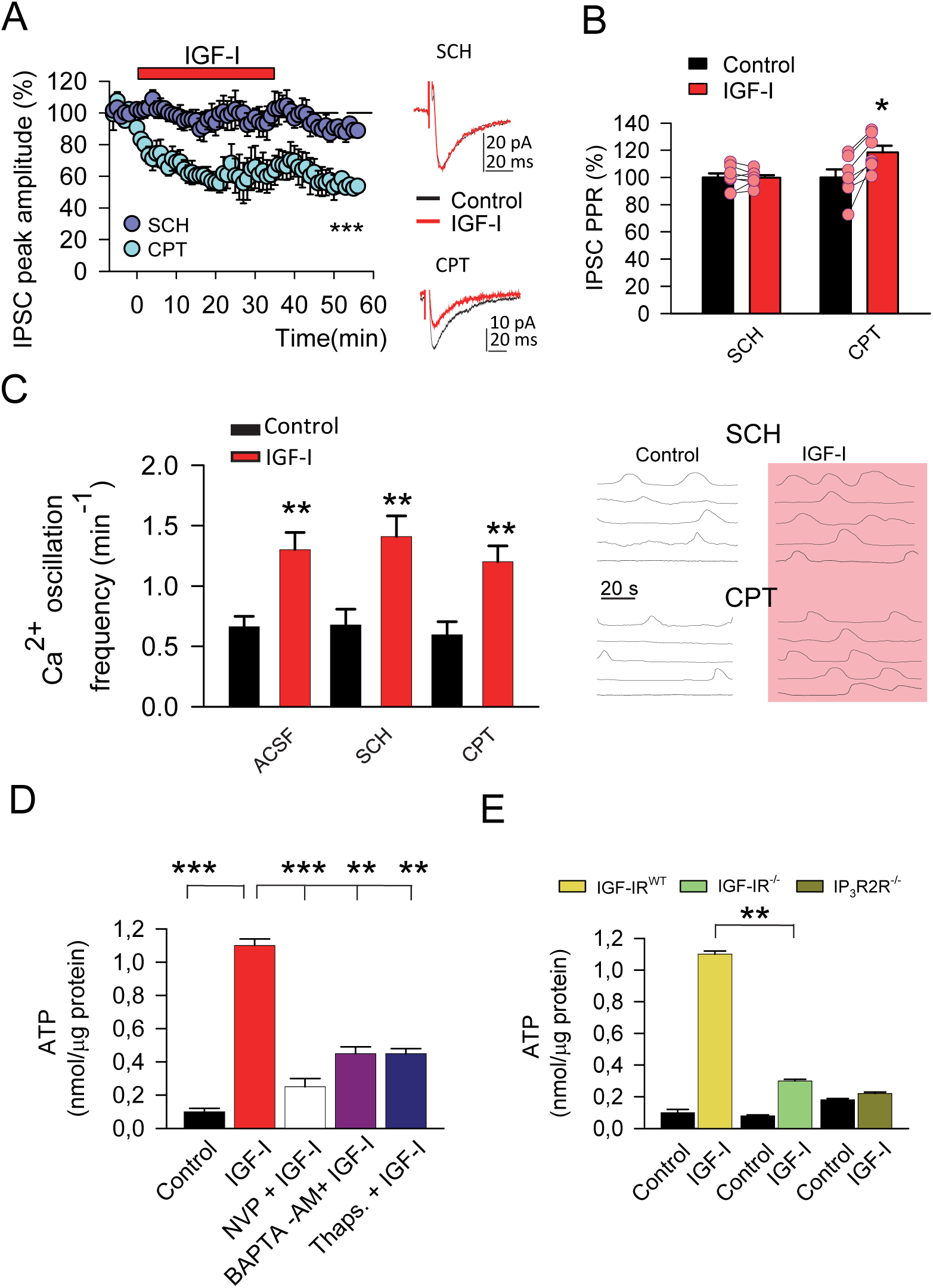
iLTD_IGFI_ requires A_2A_ receptor activation. ***A. Left.*** Same as 1B left in ACSF but in the presence of SCH or CPT (purple and blue circles respectively). Note that the A_2A_ receptor antagonist, SCH, prevents the iLTD_IGFI_. ***Right.*** Representative IPSCs in SCH and CPT before and during IGF-I. ***B.*** Same as 1D left but in the presence of SCH and CPT. ***C. Left.*** Bar plot of the calcium oscillation frequency in astrocytes in ACSF, SCH or CPT, before and during IGF-I. ***Right.*** Representative calcium traces in astrocytes under SCH or CPT, before and during IGF-I. ***D.*** Bar plot of ATP concentration in culture of astrocytes before (control) and during IGF-I exposure, and in IGF-I + NVP, IGF-I + BAPTA-AM and IGF-I + thapsigargin. ***E.*** Bar plot of ATP concentration culture of astrocytes before (control) and during IGF-I exposure from IGF-IR^WT^ (yellow bar), IGF-IR^-/-^ mice (green bar) and IP_3_R2^-/-^ mice *(gold bar)*.

### IGF-I induces ATP release from astrocytes depending on IGF-IR

We next tested whether IGF-I was capable of stimulate the release of ATP in astrocytes. We used the ATP Assay Kit (see Mat and Methods) to monitor ATP levels in cultured astrocytes before and after 1 h of treatment with IGF-I (10nM). We found that IGF-I elevated the extracellular levels of ATP (from 0.1 ± 0.02 to 1.1 ± 0.04 nmol/µg of protein before and during IGF-I, N = 6; P < 0.001; **Figure 3D**), an effect that was prevented when cultures where simultaneously treated with NVP (0.25 ± 0.05 nmol/µg of protein, during IGF-I, N = 6; P < 0.001; **Figure 3D**). The IGF-I-induced release of ATP was absent when astrocyte calcium signaling was prevented by treating cultures with BAPTA-AM (0.45 ± 0.04 nmol/µg of protein during IGF-I, N = 6; P < 0.01; **Figure 3D**). Moreover, it was also abolished by thapsigargin, which inhibits the calcium ATPase and prevents calcium mobilization from internal stores (0.45 ± 0.03 nmol/µg of protein during IGF-I, N = 6; P < 0.001; **Figure 3D**). Moreover, the release of ATP was absent when astrocytes were obtained from mice lacking IGF-IR in astrocytes (from 0.08 ± 0.005 to 0.3 ± 0.01 nmol/µg of protein, before and during IGF-I. N = 6; P < 0.01; **Figure 3E**). Finally, the release of ATP was also absent in the IP_3_R2-null mice (from 0.18 ± 0.008 to 0.22 ± 0.009 nmol/µg of protein, before and during IGF-I. N = 4; P < 0.01; **Figure 3E**), a mouse in which G-protein-mediated calcium elevation is impaired in astrocytes (42), (40). Taken together, these data suggest that IGF-I, acting through the IGF-IR in astrocytes, stimulates the calcium-dependent release of ATP.

Astrocytic activation stimulates the release of glutamate in the hippocampus, cortex and striatum (36)(41)(40),. However, iLTD_IGFI_ was unaffected by treatment with antagonists of group I metabotropic glutamate receptors (mGluRs) MPEP (50 µM) and LY367385 (100 µM; **Supplementary Figure 2**), (from 98.89 ± 0.90 to 60.78 ± 7.39 % IPSC peak amplitude, before and after IGF-I. N = 6; P < 0.01; **Figure 4A**) and the increase in the PPR induced by IGF-I was preserved (from 100.00 ± 4.06 to 120.52 ± 7.66 % of IPSC paired-pulse responses, before and after IGF-I. N = 6, P < 0.05; **Figure 4B**). Moreover, we observed that IGF-I evoked an increase in calcium event frequency in the presence of MPEP + LY367385 (from 0.51 ± 0.12 to 1.07 ± 0.14 min^-1^ before and during IGF-I. N = 44 astrocytes, P < 0.05; **Figure 4E, F**, MPEP + LY367385). Although astrocytes are able to release glutamate, these data rule out the requirement of mGLUR activation in the induction of iLTD_IGFI._

**Figure 4.**
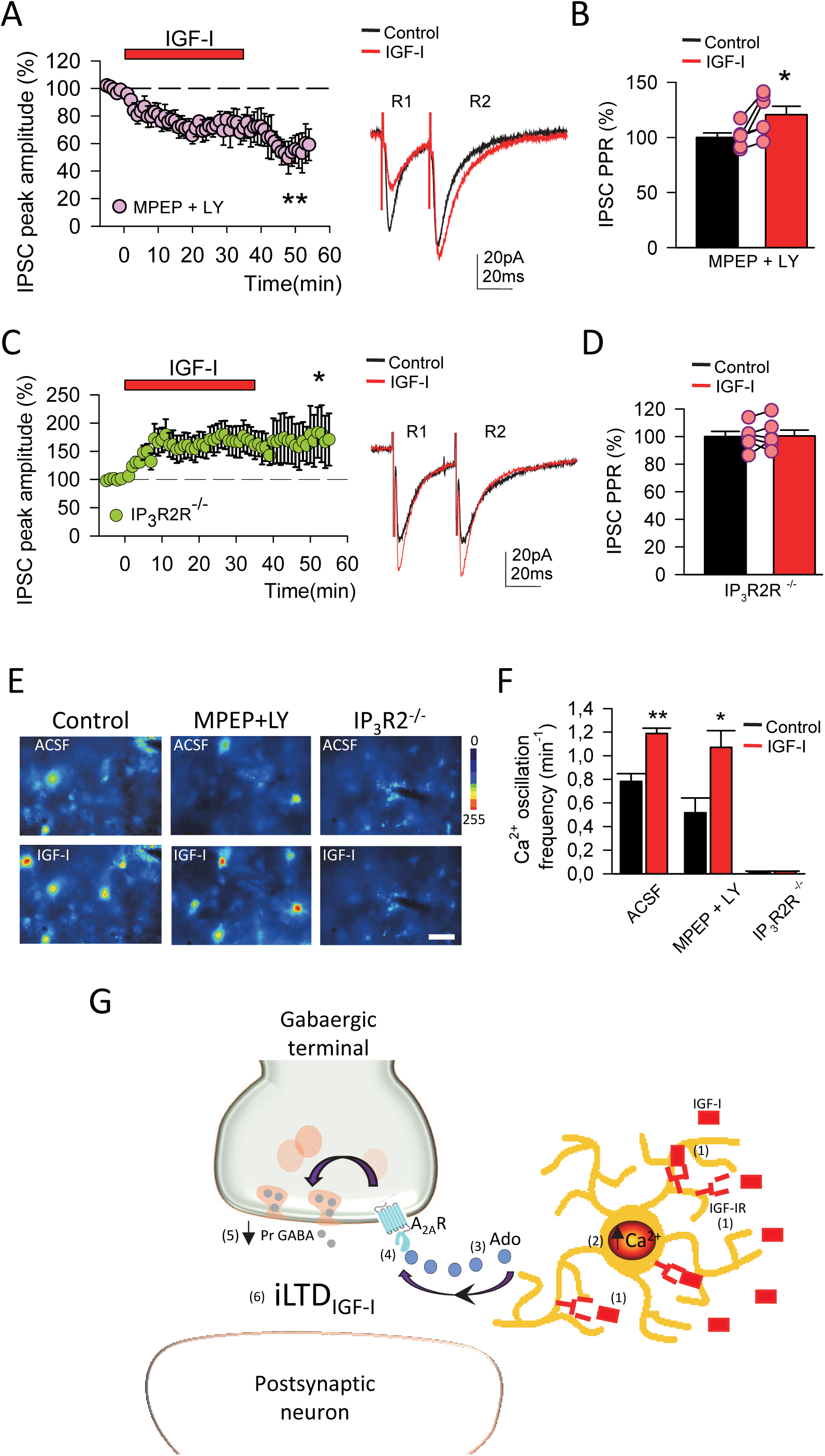
iLTD_IGFI_ is absent in IP_3_R2-null mice. ***A. Left.*** Same as 1B left in ACSF but in the presence of MPEP + LY367385. Note that in the presence of the group I mGluR antagonists, the iLTD_IGFI_ is already evoked. ***Right.*** Representative IPSCs in MPEP + LY367385 before and during IGF-I. ***B.*** Same as 1D left but in the presence of MPEP + LY367385. ***C. Left.*** Same as A left but in IP_3_R2^-/-^ mice. ***Right.*** Same as A right but in IP_3_R2^-/-^ mice. ***D.*** Same as 1D left but in IP_3_R2^-/-^ mice. ***E.*** Pseudocolor images from Fluo-4AM showing the somatic calcium elevations on astrocytes before and during IGF-I in control, MPEP + LY367385 and IP_3_R2^-/-^ mice (scale bar, 50 µm). ***F.*** Bar plot of the somatic calcium oscillation frequency in astrocytes in control, MPEP + LY367385 and IP_3_R2^-/-^ mice. ***G.*** A schematic representation of the proposed mechanism of the IGF-I-induced astrocyte-mediated LTD of the IPSCs. (1) IGF-IR activation in the astrocytes increases the astrocytic calcium frequency (2) leading to the ATP/Ado release (3) that activates presynaptic A_2A_R in the GABAergic terminal (4), decreasing the probability of GABA release (5) that underlies the LTD of the inhibitory synaptic transmission (6).

### iLTD_IGFI_ is absent in the IP_3_R2-null transgenic mice

Because cytosolic calcium elevation and ATP release from astrocytes are required for iLTD_IGF1_, we expected iLTD_IGFI_ to be absent in IP_3_R2-null mice (**Supplementary Figure 3**). The frequency of somatic calcium elevations in astrocytes of IP_3_R2^-/-^ mice was not altered during IGF-I application (from 0.001± 0.001 to 0.001 ± 0.0009 min-1 before and during IGF-I. N = 61 astrocytes, P = 1; **Figure 4E, F**, IP_3_R2^-/-^), and not only iLTD_IGFI._ was abolished, but a LTP of the IPSCs was induced (from 100.55 ± 0.64 to 167.11± 23.79 % IPSC peak amplitude, before and after IGF-I. N = 6, P < 0.05; **Figure 4C**; PPR: from 100.00 ± 3.72 to 100.39 ± 4.30 % of IPSC paired-pulse responses, before and after IGF-I. N = 6, P = 0.94; **Figure 4D**). These results further support the idea that iLTD_IGFI_ depends on IGF-IR activation in astrocytes.

### The performance of a whisker discrimination task is impaired in the astrocyte-specific IGF-IR^-/-^ mice

Finally, we tested whether the activation of IGF-IR on astrocytes was involved in the performance of a barrel cortex dependent discrimination task. We used a test based on the ability of the mice to discriminate different textures in the arms of a Y maze (**Figure 5A**). We compare the ability to perform this task on the astrocyte-specific IGF-IR^-/-^ mice (**Supplementary Figure 4A**) with their control littermates. While no differences were seen between IGF-IR^-/-^ mice and control littermates in the number of visits to the arms (**Figure 5B**), indicating normal deambulatory activity, IGF-IR^-/-^ mice spent less time examining the arm with the novel texture (N = 11; P = 0,0336; **Figure 5C left**), indicating impaired texture discrimination (**Figure 5C right**). Conversely, whisker perception in IGF-IR^-/-^ mice was preserved, as indicated by normal performance in the gap-crossing test (**Supplementary Figure 4B**). In addition, working memory was also normal in IGF-IR^-/-^ mice, as indicated by preserved performance in the Y maze alternation test (**Supplementary Figure 4C**). Taken together these results indicate that IGF-IR on astrocytes is involved in the texture discrimination in mice.

**Figure 5.**
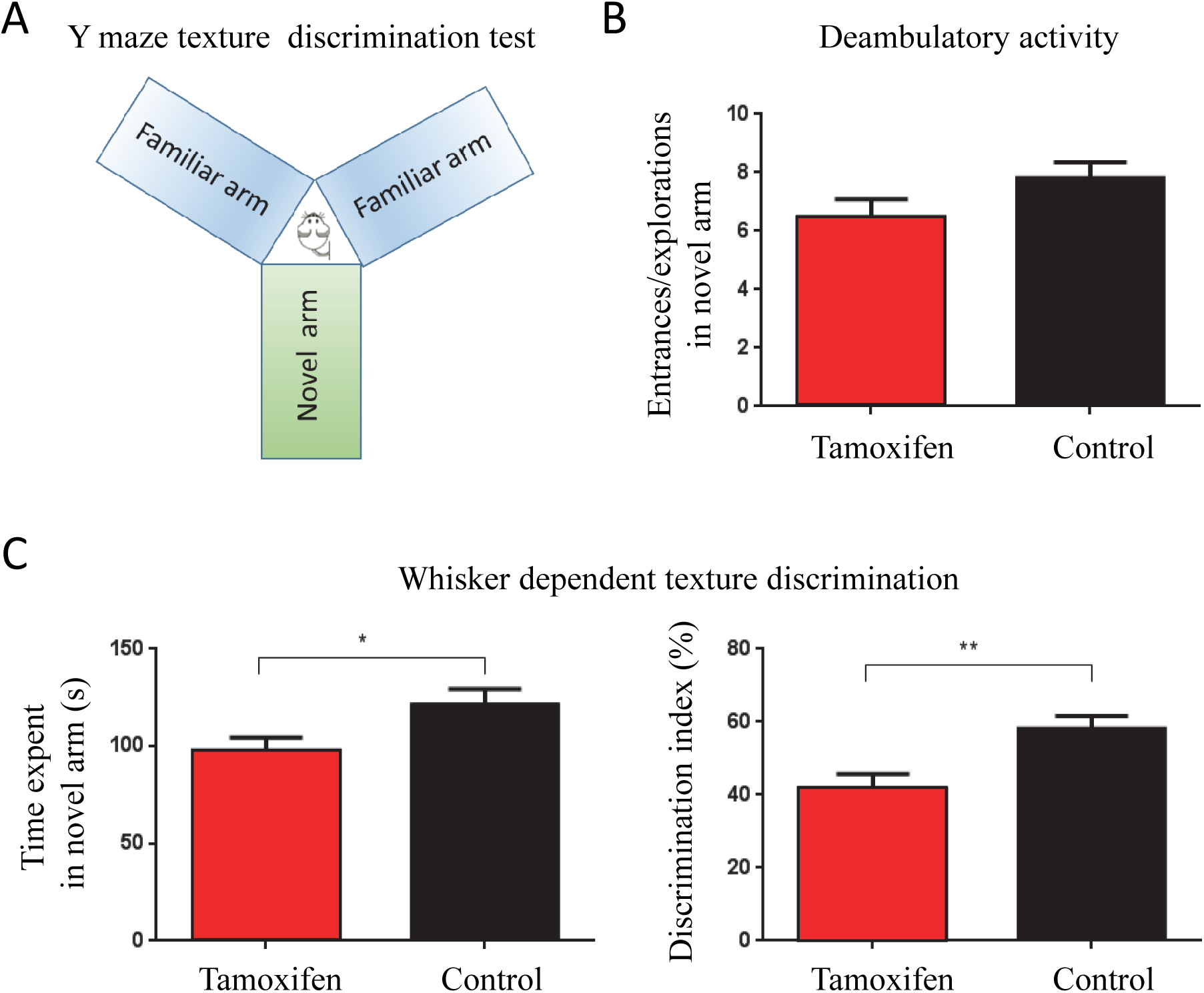
Impaired whisker-based discrimination in IGF-IR KO mice. ***A.*** Schema of Y maze whisker dependent texture discrimination test. While two arms (familiar) were covered with a 500 grit sandpaper, the third (novel) was covered with one of 220 grit. Since the three arms of the maze are identical, and there are no extra-maze cues, discrimination of novelty vs familiarity relies only on the different textures that the animal can perceive with the whiskers. **B.** IGF-IR^-/-^ and control mice show no differences in entrance to the different arms (p=0.1142). ***C left.*** However, control mice spent more time exploring the novel arm (220 grift sandpaper) than IGF-IR^-/-^ mice (n=13, *p=0,0336). ***C Right.*** Percentage of time spent in the novel-texture arm was higher in controls than in IGF-IR^-/-^ mice (**p=0,0059; unpaired student t-test with Welch’s correction; control =11, IGF-IR^-/-^ =13).

## DISCUSSION

The operation of cortical circuits depends on a fast time course of inhibitory signaling in interneurons, whereas modulation of synaptic inhibition plays a crucial role in the induction of cortical plasticity. Our results show that IGF-I can induce a long lasting depression of the IPSCs that we have labeled as iLTD_IGFI_, by activating IGF-IRs in astrocytes. The latter induces a calcium-dependent release of ATP/Ado, and iLTD_IGFI_ depends on the activation of A_2A_ adenosine receptors (see mechanistic cartoon). These presynaptic adenosine receptors would induce a decrease in the probability of GABA release at inhibitory terminals and an increase in PPRs that parallels iLTD_IGFI_. The lack of iLTD_IGFI_ in IP_3_R2^-/-^ mice and in mice in which IGF-IR has been selectively ablated in astrocytes, demonstrates that both IGF-IR activation in astrocytes and somatic cytosolic calcium elevations are crucial in the induction of this novel form of long lasting down regulation of cortical synaptic inhibition. Although IP_3_R2^-/-^ mice present residual Ca^2+^ events that might occur in fine processes(43), it seems that the existence of those events would not be sufficient to induce iLTD_IGFI_. Quite the opposite, an intriguing LTP of the IPSCs is revealed. Thus, these data suggest that although astrocyte-synaptic interactions might take place at astrocyte processes, iLTD_IGFI_ requires an active contribution of astrocytic somatic Ca^2+^ signaling. Importantly, IP_3_R2^-/-^ mice have recently been reported to show impaired LTD and memory deficits (44).

IGF-I elicits a long-lasting depression of GABA release by cerebellar Purkinje cells in response to glutamate, indicating that IGF-I may act as a modulator of glutamatergic transmission in the adult rat olivo-cerebellar system (10)(45). Also, IGF-I has been shown to modulate GABAergic transmission in the olfactory bulb (13). However, an IGF-I mediated long lasting depression of fast GABAergic synaptic transmission in the neocortex has not been described before. Here we demonstrate that IGF-I induces the release of ATP from cortical astrocytes, that in turn is converted to adenosine to induce long-lasting depression of GABA release. Although IGF-IRs may be present in inhibitory GABAergic terminals, there are no clear data showing a direct action of IGF-I acting on presynaptic IGF-IRs. The only evidence suggesting this mode of action is seen in the hippocampus where IGF-I, acting possibly via GABAergic neurons, can induce the release of GABA to regulate endogenous ACh release (11). At any rate, our observations contribute to the notion that IGF-I can modulate both stimulatory and inhibitory synaptic activity throughout the CNS.

In rodent astrocytes, IGF-I signaling is important for glucose uptake, regulation of glutamate transport and protection against oxidative stress in the brain (46)(47)(48). Indeed, loss of IGF-IR has been associated with increased GLUT1 activity and, consequently, increased glucose uptake (25). Although there is evidence showing that insulin signaling in astrocytes mediates tyrosine phosphorylation of Munc18cand syntaxin-4–dependent ATP exocytosis, which in turn modulates presynaptic dopamine release (49), we have not found previous evidence for an stimulatory action of IGF-I on ATP release by astrocytes. In this regard, it has been previously shown that metabotropic Protease-activated receptor 1 (PAR-1) induces exocytosis of ATP from cortical astrocytes, which leads to a short term downregulation of inhibitory synaptic currents in layer 2/3 pyramidal neurons (33). In contrast with the presynaptic iLTD described herein, this short term modulation is mediated by a postsynaptic mechanism in which Ca^2+^-entry through the neuronal P2X purine receptor leads to a phosphorylation-dependent down-regulation of GABA_A_ receptors. Therefore, astrocytes can down- or up-regulate inhibitory synaptic transmission by a calcium-dependent release of ATP depending on whether IGF-IR or PAR1R are activated, respectively.

Astrocytes have been shown to respond to both glutamate and GABA (28)(50)(51)(36)(52), which allows them to sense the activity of excitatory and inhibitory neurons. In response to these neurotransmitters they can release both glutamate and ATP/Ado (53)(35)(54)(55)(56). Indeed, hippocampal interneuron activity leads to GABA_B_R-mediated release of glutamate from astrocytes that potentiates both inhibitory (28), and excitatory (29) synaptic transmission. In addition to glutamate, hippocampal astrocytes may also release ATP, which is converted to adenosine that depress (57)(30)(31)(32) or potentiates excitatory synaptic transmission (38). Moreover, ATP released by astrocytes can depress excitatory synapses from basolateral amygdala and enhance inhibitory synapses from the lateral subdivision of the central amygdala via the activation of A_1_ and A_2A_ adenosine receptors, respectively (58). Furthermore, cortical astrocytes have been shown to induce a short-term depression of inhibitory synaptic currents (33). However, to our knowledge, we present the first evidence showing that IGF-IR activation in astrocytes can induce a long lasting depression of inhibitory cortical synaptic transmission through the release of ATP/Ado via a presynaptic mechanism.

At the circuit level, interneurons control the flow of information and synchronization in the cerebral cortex. Synaptic inhibition is involved in the emergence of fast brain rhythms (59), and in the induction of synaptic plasticity (60), that jointly contribute to cognitive functions. Indeed, disruption of astrocytic vesicular release has been found crucial for gamma oscillatory hippocampal activity with significant impact in recognition memory tasks (61). Our results demonstrate that IGF-I receptor on astrocytes improves the performance of the texture discrimination in mice, suggesting that the iLTD_IGFI_ could be essential in this barrel cortex dependent task. The release of ATP/Ado from astrocytes and the iLTD_IGFI_ described here could play a relevant role in this cognitive function by controlling brain rhythms and favoring the induction of Hebbian synaptic plasticity. Decreased synaptic inhibition would facilitate the back-propagation of action potentials into the dendrites and the induction of spike timing-dependent plasticity. In addition, by changing the ratio between synaptic excitation and inhibition, neuronal membranes can rapidly reach the threshold for action-potential generation, and enhanced cortical activity is expected when cortical levels of IGF-I increase. Indeed, this increase in cortical activity would activate neurotrophic coupling mechanism of entrance of IGF-I from the plasma into the brain (2).

In summary, the present findings reveal novel mechanisms and functional consequences of IGF-I signaling in the cortex. It induces the long-term depression of GABAergic inhibition (iLTD_IGFI_) and regulates the behavioral performance in the barrel cortex-related texture discrimination tasks, though activation of cortical astrocytes.

## METHODS

### Materials

2-(2-Furanyl)-7-(2-phenylethyl)-7*H*-pyrazolo[4,3-*e*][1,2,4] triazolo [1,5-*c*] pyrimidin-5-amine (SCH58261 (SCH)), 8-Cyclopentyltheophylline (CPT) and thapsigargin were purchased from Tocris. NVP AEW541 (NVP) was purchased from Cayman. Recombinant Human IGF-I (IGF-I) was from Peprotech. BAPTA-K4 was from Santa Cruz Byotechnology. Fluo-4-AM was purchased from Invitrogen and ATP Assay Kit was from Abcam. The rest of drugs were purchased from Sigma-Aldrich.

### Ethics statement and animals

All animal procedures were approved by the Ethical Committee of the Universidad Autónoma of Madrid, and Cajal Institute and are in accordance with Spanish (R.D. 1201/2005) and European Community Directives (86/609/EEC and 2003/65/EC), which promote animal welfare. Male C57BL/6J or transgenic mice were housed under a 12-h/12-h light/dark cycle with up to five animals per cage and were used for slice electrophysiology. Transgenic mice with a deletion of IGF-IR in astrocytes (GFAP-IGF-IR^-/-^ mice) were obtained by injecting IGF-IR^f/f^ mice (B6, 129 background; Jackson Labs) with AAV8-GFAP-mCherry (UMN vector core) or AAV8-GFAP-mCherry-CRE viral vectors (UNC vector core). For a subset of experiments, slices were obtained from male IP_3_R2^-/-^ mice generously donated by Gertrudis Perea (Cajal Institute).

### Slice preparation

Wild type and IP_3_R2^-/-^ mice (12-18 days old), as well as GFAP-IGF-IR^-/-^ mice and their controls (12-20 weeks of age) were sacrificed, and brains submerged in cold (4°C) cutting solution containing (in mM): 189.0 sucrose, 10.0 glucose, 26.0 NaHCO_3,_ 3.0 KCl, 5.0 Mg_2_SO_4_, 0.1 CaCl_2_, 1.25 NaH_2_PO_4_.2H_2_O. Coronal slices (350 µm thick) were cut with a Vibratome (Leica VT 1200S), and incubated (>1h, 25–27°C) in artificial cerebrospinal fluid (ACSF) containing (in mM): 124.00 NaCl, 2.69 KCl, 1.25 KH_2_PO_4_, 2.00 Mg_2_SO_4_, 26.00 NaHCO_3_, 2.00 CaCl_2_, and 10.00 glucose). pH was stabilized at 7.3 with a 95 % O_2_, 5 % CO_2_ mixture. Slices were transferred to a 2 ml chamber fixed to an upright microscope stage (BX51WI; Olympus, Tokyo, Japan) equipped with infrared differential interference contrast video (DIC) microscopy and superfused at room temperature with 95 % O_2_, 5 % CO_2_-bubbled ACSF (2 ml/min).

### Electrophysiological recordings

Patch-clamp recordings from layer 2/3 pyramidal neurons of the barrel cortex were performed in whole-cell voltage-clamp configurations with patch pipettes (4–8 MΩ) filled with an internal solution that contained (in mM): 135 KMeSO_4_, 10 KCl, 10 HEPES-K, 5 NaCl, 2.5 ATP-Mg^+2^, and 0.3 GTP-Na^+^, pH 7.3. In some experiments, a BAPTA-based intracellular solution was used (in mM): 40 BAPTA-K_4_, 2 ATP-Na^2+^, 10 mM HEPES, 1 MgCl_2_ and 8 NaCl, pH 7.3 Recordings were performed using a Cornerstone PC-ONE amplifier (DAGAN, Minneapolis, MN). Pipettes were placed with a mechanical micromanipulator (Narishige, Tokyo, Japan). The holding potential was adjusted in a range from −70 to −80 mV, and series resistance was compensated to ≈ 80 %. Layer 2/3 pyramidal neurons located over the barrels (layer 4) were accepted only when the seal resistance was >1 GΩ and the series resistance (10–20 MΩ) did not change (>20 %) during the experiment. Data were low-pass filtered at 3.0 kHz and sampled at 10.0 kHz, through a Digidata 1550B (Molecular Devices, Sunnyvale, CA).

### Synaptic stimulation

Inhibitory synaptic currents (IPSCs) were isolated in the presence of AMPAR and NMDAR antagonists (20 µM CNQX and 50 µM D-AP5, respectively). IPSCs were evoked with a bipolar stimulation electrode pulled from theta borosilicate glass capillary (diameter of the tip, 5–8µm), filled with ACSF, and connected to a Grass S88 stimulator and a stimulus isolation unit (Quincy, USA digital stimulator) through chloride-silver electrodes. The stimulating electrodes were placed at layer 4 of the barrel cortex near the tip (∼100-150 µm) of the recording pipette. Paired pulses (200-μs duration and 50 ms-interval) were continuously delivered at 0.33 Hz. After recording 5 minutes of stable baseline of IPSCs, IGF-I was added to the bath during 35 minutes. After that, IGF-I was washed for at least 10 minutes and the amplitude of the IPSCs after IGF-I washout was continuously checked. The pre- or postsynaptic origin of the observed regulation of synaptic currents was tested by estimating changes in the paired pulse ratio (PPR) of the IPSCs (62)(63)(64). We calculated the PPR index (R2/R1), where R1 and R2 are the peak amplitudes of the first and second IPSC, respectively.

### Stereotaxic surgery and virus delivery

Animals between 6-10 weeks’ old were anesthetized with ketamine (100 mg/kg)/xylazine (10 mg/kg) mix. To delete IGF-Irs selectively in cortical astrocytes, IGF-IR^f/f^ mice were injected in the barrel cortex with 500 nl of the viral vector AAV8-GFAP-mCherry-CRE (or AAV8-GFAP-mCherry as a control). To monitor astrocyte calcium levels, mice were injected with AAV5-GFAPABC1D-cytoGCaMP6f-SV40 (Penn Vector Core). Injections were made at layer 2/3 of barrel cortex with a Hamilton syringe attached to a 29-gauge needle at a rate of 0.5 μl/ min. Coordinates used to reach the area were: anterior–posterior: −1.0 mm; medial–lateral: +/−3.50 mm; dorsal–ventral: −0.15 mm. Three weeks after the injection, successful delivery was confirmed by the location of the virus based on mCherry expression.

### Ca^2+^ imaging

In GFAP-IGF-IR^-/-^ mice and their controls, Ca^2+^ levels in astrocyte soma and processes were obtained by two-photon microscopy (Leica DM6000 CFS upright multiphoton microscope with TCS SP5 MP laser) using the GCaMP6f viral vector targeted to astrocytes. In some experiments, somatic Ca^2+^ levels were obtained by using epifluorescence microscopy and the Ca^2+^ indicator Fluor-4-AM (5 μM in 0.01% of pluronic, 30-45 min incubation at room temperature). Loaded cells were illuminated every 100 ms at 490 nm with a monochromator (Polychrome IV; TILL Photonics), and successive images were obtained at 1 Hz with a cooled monochrome CCD camera (Hammamatsu, Japan) attached to the Olympus microscope that was equipped with a filter cube (Chroma Technology). Camera control and synchronization for the epifluorescence were made by ImagingWorkbench software (INDEC-BioSystems) and for two-photon imaging by the Leica LAS software.

### Astrocyte cultures and ATP release

Pure astroglial cultures were prepared as described(65). Postnatal (day 0-2) brains from C57BL/6J (WT) and constitutively GFAP IGF-IR^-/-^ mice(25) were removed and immersed in ice-cold Earle’s balanced salt solution (ThermoFisher, Waltham, MA USA). Cortices were dissected and mechanically dissociated. The resulting cell suspension was centrifuged and plated in DMEM/F-12 (ThermoFisher) with 10% fetal bovine serum (FBS, ThermoFisher) and 100mg/ml of antibiotic-antimycotic solution (Sigma, St. Louis, MO, USA). Cell cultures were maintained in an incubator at 37 °C in 95% humidity with 5% CO2. Pure astrocytes monolayer cultures were replated at 2.5 × 10^5^ cells/well in DMEM/F-12 with 10% FBS medium. After two days, medium was replaced by DMEM/F-12 during 3 hours followed by one-hour treatment with 10 nM IGF-I (PreproTech, Rocky Hill, NJ, USA) or one-hour treatment with inhibitors: 400 nM NVP-AEW541 (IGF-I receptor antagonist, Cayman, Ann Arbor, MI, USA), 10 µM BAPTA-AM (intercellular Ca2+ chelator, Sigma) or 1 µM Thapsigargin (endoplasmic reticular Ca2+-ATPase inhibitor, Tocris, Bristol, UK) followed by one-hour stimulation with 10 nM IGF-I.

The amount of ATP released into the medium was measured with an ATP Assay Kit (Fluorometric, ab83355; Abcam, Cambridge, UK) according to the manufacturer’s instructions. Firstly, cells were washed with cold phosphate-buffered saline and resuspended in 100 µl of ATP assay buffer. Then, the cells were quickly homogenized by pipetting up and down, and centrifuged for 2 min at 4°C (18 000 g) to remove any insoluble material. Thereafter, the supernatant was collected and transferred to a clean tube and kept on ice. ATP assay buffer (400 µL) and 100µL of ice cold 4M perchloric acid were added to the homogenate to deproteinate the samples. Homogenates were vortexed briefly and incubated on ice for 5 min, centrifuged at 18 000 g for 2 min at 4°C, and the supernatant transferred to a fresh tube. The supernatant volume was measured and an equal volume of ice cold 2 M KOH was added. Finally, the homogenate was centrifuged at 18 000 g for 15 min at 4°C and the supernatant collected. ATP reaction mix (50 µL) and 50 µL of sample were added to each well and incubated at 24 °C for 30 min, protected from light. Samples were measured on a FLUOstar microplate reader (BMG LabTech, Ortenberg, Germany) at 535/587 nm. Each sample was run in duplicate. The concentrations of ATP released from astrocyte primary cultures were obtained from standard curves and normalized to the total amount of protein.

### Behavioral experiments

Adult male and female control (C57BL/6JolaHsd; 6-12 months old; 28-35g), and transgenic mice were used in these studies. Transgenic mice with tamoxifen-regulated deletion of IGF-IR in astrocytes (IGF-IR^-/-^ mice) were obtained by crossing IGF-IRf/f mice (B6, 129 background (IGF-IRflox/flox mice; Jackson Labs; stock number: 012251) with CreERT2.GFAP mice (C57B&/6xSJL/J mix background; Jackson Labs, stock number: 012849, see (66) for further details). In these animals, IGF-IR activity is knocked down after administration of tamoxifen, as described elsewhere (67). Tamoxifen was injected for 5 consecutive days intraperitoneally (75 mg/kg, Sigma) at the age of 1 month, and animals were used at 2-7 months old. Controls used were siblings treated with the vehicle for tamoxifen injections (corn oil). Using the tdTomato/eGFP reporter mouse to detect Cre-mediated deletion in response to tamoxifen administration in CreERT2.GFAP mice, we previously documented that it was restricted to astrocytes (68). Multiplex PCR for mouse genotyping included a common forward primer (P3, 5’−CTG TTT ACC ATG GCT GAG ATC TC−3’) and two reverse primers specific for the wild-type (P4, 5’−CCA AGG ATA TAA CAG ACA CCA TT−3’) and mutant (P2, 5’-CGC CTC CCC TAC CCG GTA GAA TTC-3’) alleles. GFAP-T-IGF-IR mice show lower levels of IGF-IR in brain after tamoxifen injection. Mice were housed in standard cages (48 × 26 cm2) with 4 - 5 animals per cage, kept in a room with controlled temperature (22°C) under a 12-12h light-dark cycle, and fed with a pellet rodent diet and water ad libitum. All experimental protocols were performed during the light cycle between 14 pm - 18 pm.

Animal procedures followed European guidelines (2010/63/EU) and were approved by the local Bioethics Committee (Madrid Government). *qPCR:* Total RNA isolation from brain tissue was carried out with Trizol. One mg of RNA was reverse transcribed using High Capacity cDNA Reverse Transcription Kit (ThermoFisher Scientific, USA) according to the manufacturer’s instructions. cDNA (62.5 ng) was amplified using TaqMan probes for mouse IGF-I receptor (IGF-IR), and rRNA 18S as endogenous control (ThermoFisher Scientific). Each sample was run in triplicate in 20 µl of reaction volume using TaqMan Universal PCR Master Mix according to the manufacturer’s instructions (ThermoFisher Scientific). All reactions were performed in a 7500 Real Time PCR system (ThermoFisher Scientific). Quantitative real time PCR analysis was carried out as described (69). Results were expressed as relative expression ratios on the basis of group means for target transcripts versus reference 18S transcript.

For behavioral tests, both sexes were used and balanced between experimental groups. Since wild type and control littermates performed similarly in the tests, they were pooled and presented as a single control group. We performed the following three behavioral tests:

1. *Gap-crossing test*. To assess sensory perception through the whiskers we used the gap-crossing test that consists of a series of trials in which the mouse has to cross a gap with increasing distances (70)(71). Each mouse was placed in the center of an elevated platform (5 cm wide, 9 cm long) connected to a safe black cylindrical tube (8 cm diameter, 9 cm long). Gap distance between the platform and the cylindrical tube ranged from 0 to 8 cm in 1 cm increments. The test was performed under infrared lighting, the trials were recorded with a video camera and the maximum distance crossed by each animal was measured.
2. *Y-maze spontaneous alternation test*. Working memory was assessed by recording spontaneous activity while exploring a Y-maze (72). The maze was made of black-painted wood and each arm was 25 cm long, 14 cm high, 5 cm wide, and positioned at equal angles. Trials lasted 8 min each. After each trial, the maze was cleaned with 70% ethanol to remove olfactory cues. Offline analysis of the videos was carried out to obtain the sequence of entries during the test. Alternate behavior was calculated as the percentage of real alternations (number of triplets with non-repeated entries) versus total alternation opportunities (total number of triplets).
3. *Whisker discrimination test*. To assess the ability of the animals to discriminate different textures with their whiskers, we adapted the two-trial Y-maze test described in previous work (73)(74). The apparatus was constructed in black-painted wood with three arms, each 25 cm long, 5 cm wide, and 14 cm high. The walls of the maze arms were covered with two different grades of black sandpaper. While two arms (familiar) were covered with a 500 grit sandpaper, the third (novel) was covered with one of 220 grit. Since the three arms of the maze are identical, and there are no extra-maze cues, discrimination of novelty vs familiarity relies only on the different textures that the animal can perceive with the whiskers. Experiments were conducted in a room with dim illumination (6 lux). During the acquisition phase, mice were placed at the end of one of the familiar arms (in a random order) and were allowed to explore both familiar arms (500 grit sandpaper) for 5 min while the third arm (novel; 220 grit sandpaper) was closed with a guillotine door. At the end of the first trial, mice were returned to their home cage for 5 min. In the retrieval phase, the mice were placed again at the end of the same arm where they started the acquisition phase, and allowed to freely explore all three arms for 5 min. To remove possible olfactory cues, the maze was cleaned with 70% ethanol between the trials. The time spent in each of the arms was recorded using a video camera, and the discrimination index [novel arm/ (novel+familiar arms)] × 100 calculated.

### Data analysis

Electrophysiological data analysis was done in Clampfit 10 (Axon Instrument) and ImageJ (NIH, Bethesda, Maryland, USA) was used for the calcium imaging. Graphs were drawn in SigmaPlot 11 (Systat Software Inc, San Jose, CA, USA). Fluorescence values are given as ΔF/F_0_ (ΔF/F_0_= 100 x (F-F_0_)/F_0)_, where F_0_ is the pre-stimulus fluorescence level when cells were at rest, and F is the fluorescence at different times during activity. We used the Student’s two-tailed *t*-tests for unpaired or paired data as required in the electrophysiology experiments. Data are presented as means ± SEM. The threshold for statistical significance was *P*< 0.05(*); *P*< 0.01 (**) and *P*< 0.001 (***) for the Student’s test. Statistical analysis was performed using GraphPad Prism 5.0 (La Jolla, CA, USA). To compare differences between two groups and compare multiple variables, two-way ANOVA was used, followed by a Bonferroni post hoc test to compare replicate means. Statistical differences were considered when p < 0.05. Results are presented as mean ± SEM of five independent experiments. (*p< 0.05, **p < 0.01, ***p < 0.001).

## ACKNOWLEDGMENTS

This work was supported by the following Grants: BFU2016-80802-P AEI/FEDER, UE (MINECO) to D. Fernández de Sevilla, National Institutes of Health-NINDS (R01NS097312 and R01DA048822) to A. Araque and grants from Ciberned and SAF2016-76462-C2-1-P (MINECO) to I. Torres-Aleman. JAZV acknowledges the support from the National Council of Science, Technology and Technological Innovation (CONCYTEC, Perú) through the National Fund for Scientific and Technological Development (FONDECYT, Perú). JF received a post-doc fellowship from Fundação de Amparo à Pesquisa do Estado de São Paulo (FAPESP: # 2017/14742-0; # 2019/03368-5). We thank the University of Minnesota Viral Vector and Cloning Core for production of some of the viral vectors used in this study. The authors thank Dr. J. Chen (UCSD, CA, USA) for providing IP_3_R2^-/-^ mice and Dra G. Perea for helpful comments.

## AUTHOR CONTRIBUTIONS

Noriega-Prieto and Fernandez de Sevilla conceived the study. Noriega Prieto, Maglio performed and analyzed electrophysiological, and calcium imaging experiments, analyzed the data and wrote the methods. Jonathan and Jansen designed and performed behavioral tests and wrote the methods. Martinez-Rachadell performed behavioral tests. Fernandez performed the in vitro tests with astrocytes, characterized IGF-IR^-/-^ mice and wrote the methods and results. Pignatelli prepared and validated the virus and the IGF-IR^-/-^ mice. Torres-Aleman, Araque, Nuñez and Fernandez de Sevilla designed, coordinated, interpreted experiments and wrote part of the manuscript. All the authors read and edited the manuscript.

## SUPPLEMENTARY INFORMATION

**Supplementary Figure 1.**
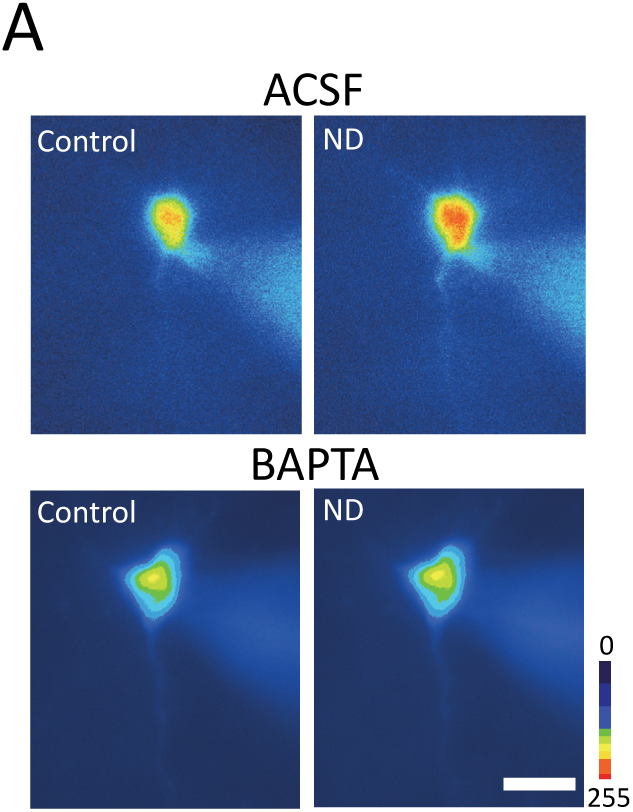
Elevations in astrocytes somatic calcium are absent in IP_3_R2 null mice. ***A.*** Pseudocolor images showing the astrocyte calcium increase during ATP (100 µM) bath perfusion in wild type (upper images) and IP_3_R2^-/-^ (bottom images) mice (scale bar, 50 µm).

**Supplementary Figure 2.**
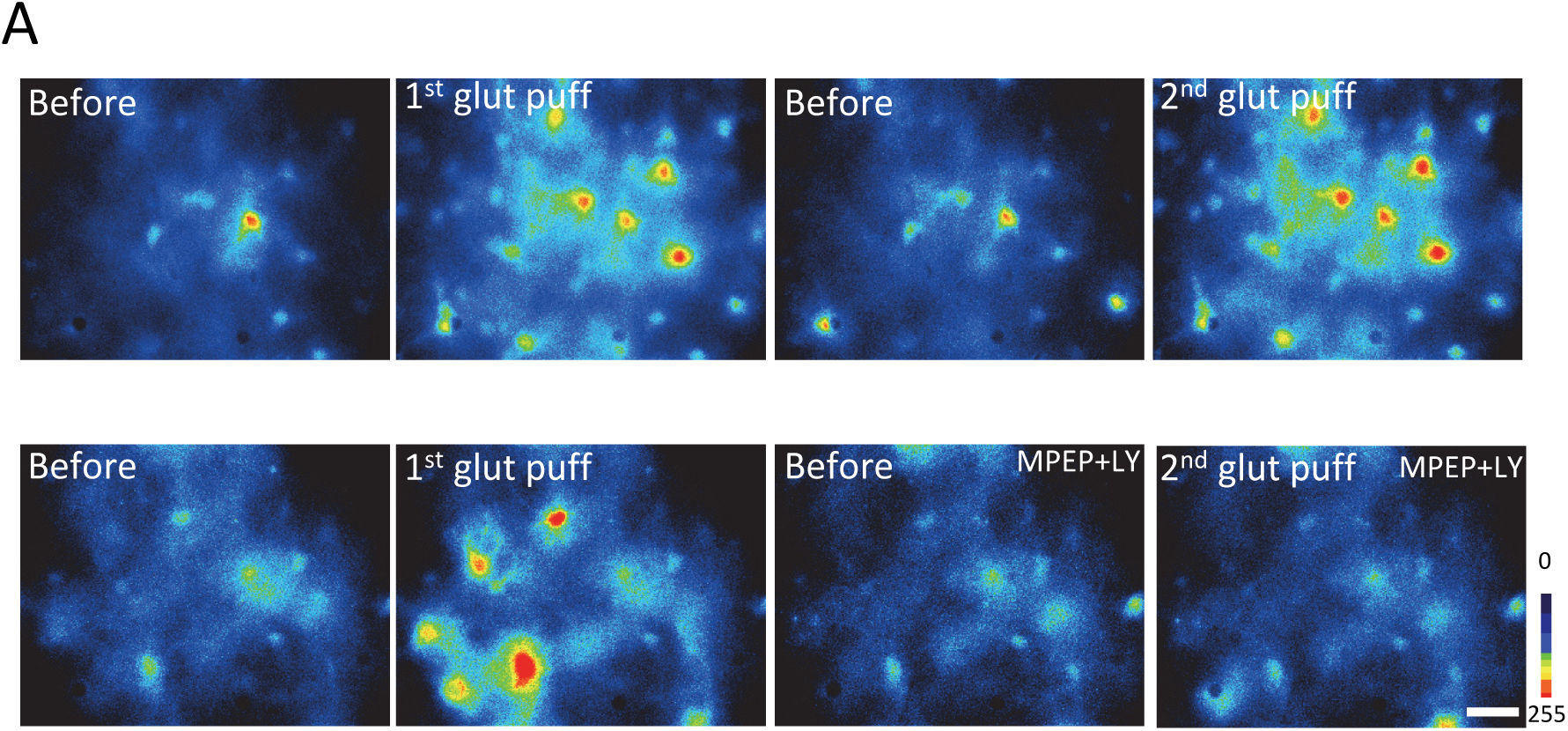
Postsynaptic BAPTA blocks the intracellular calcium elevations. ***A.*** Pseudocolor images of recorded cells before and during neuronal depolarization (ND) in control (ACSF, upper images) and with intracellular BAPTA (BAPTA, bottom images; scale bar, 50 µm).

**Supplementary Figure 3.**
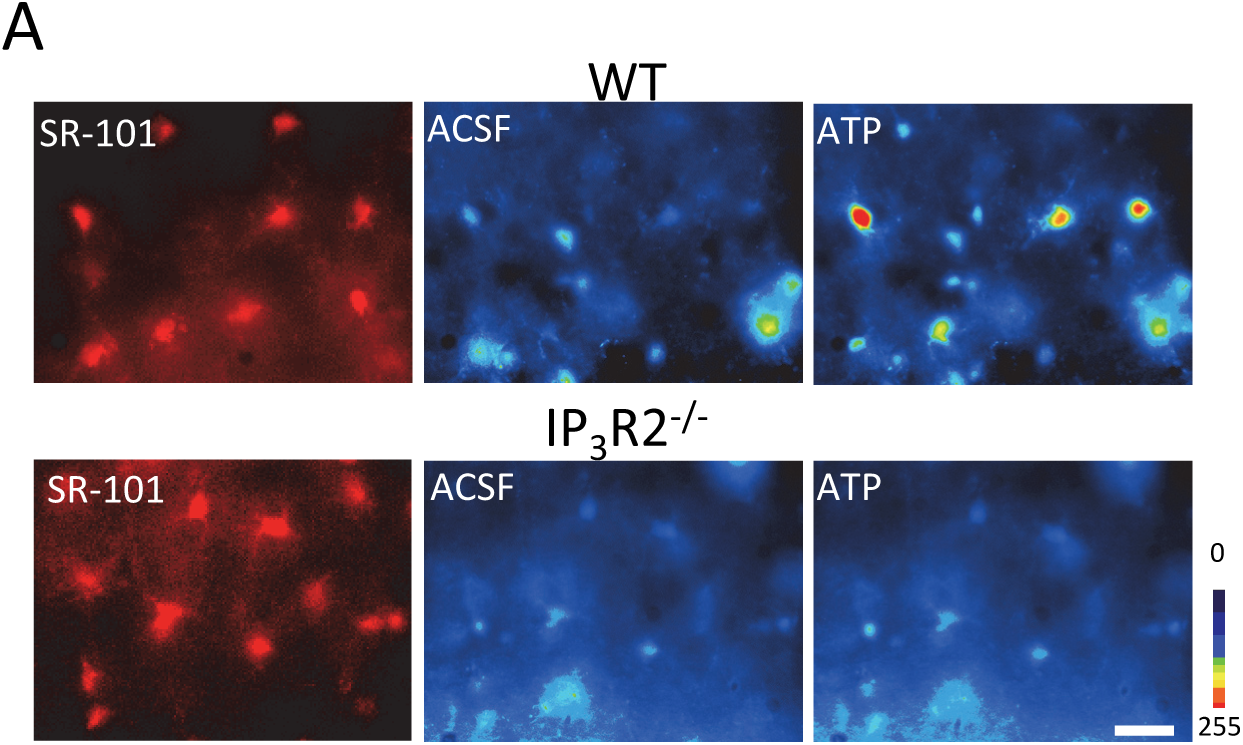
Group I mGluR antagonists decrease calcium elevations in astrocytes. ***A.*** Pseudocolor images of Fluo 4AM-filled astrocytes in the presence of TTX (1µM), CNQX (20 µM) and D-AP5 (50 µM), before and during two consecutive glutamate (10 mM) pressure pulses (2 s-duration,) in control (upper images) and during MPEP + LY367385 (bottom images) bath perfusion (scale bar, 50 µm).

**Supplementary Figure 4.**
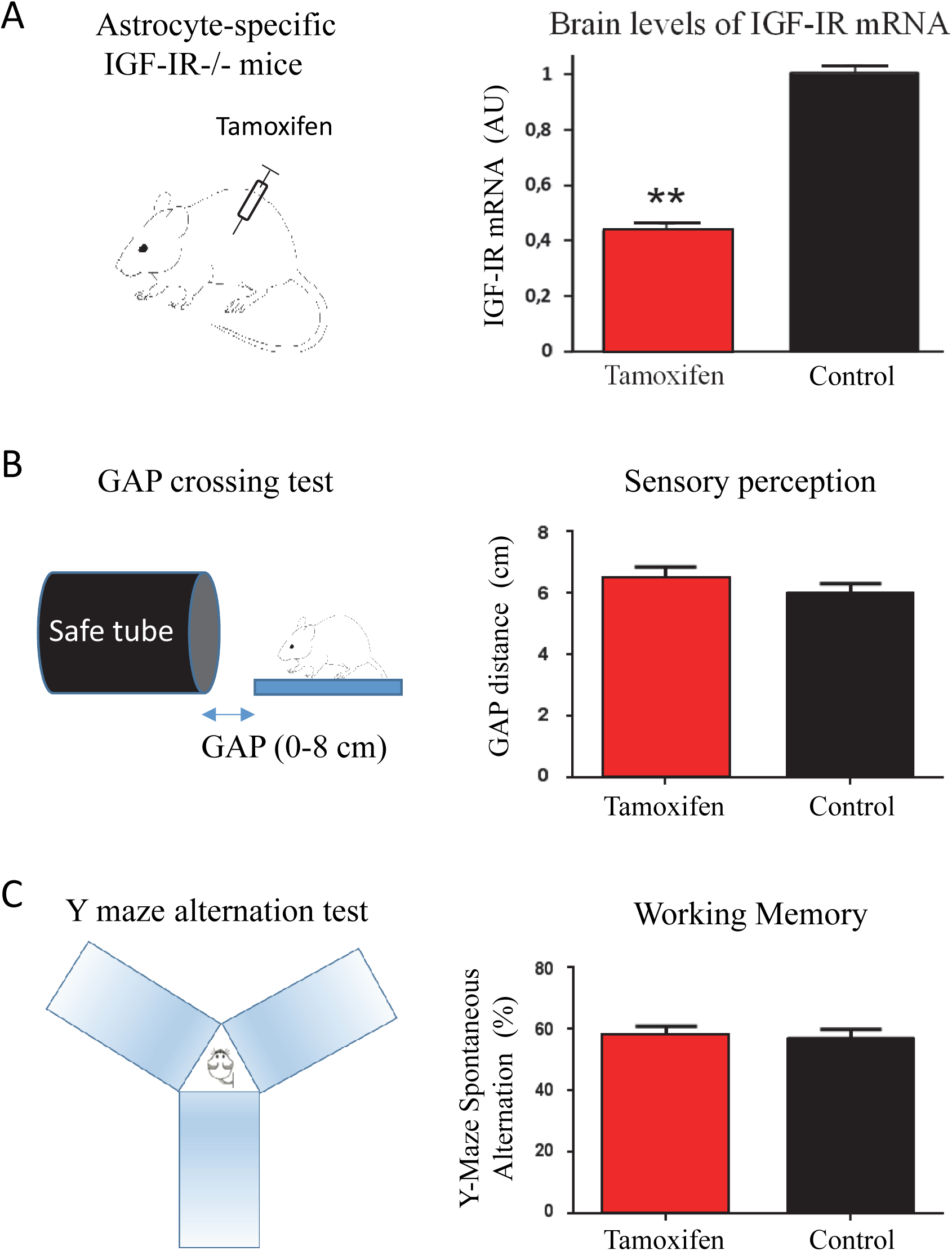
Preserved sensory perception and working memory in IGF-IR KO mice. ***A left.*** Schema of the transgenic mice generation with tamoxifen-regulated deletion of IGF-IR in astrocytes (IGF-IR^-/-^ mice). ***A right***, Brain levels of IGF-IR mRNA, as determined by qPCR, are reduced in IGF-IR^-/-^ mice after tamoxifen injection, as compared to vehicle-injected control mice (**p<0.01; n=5 per group). ***B left,*** Schema of the GAP crossing test used to evaluate the sensory perception through the whiskers. The gap-crossing test consists of a series of trials in which the mouse has to cross a gap with increasing distances (0 to 8 cm). ***B right,*** Control and IGF-IR^-/-^ mice showed similar performance (control=5, IGF-IR^-/-^ =7, p=0,6199) in the Gap-crossing test, indicating no alterations in the sensory perception. ***C left,*** Schema of Y maze spontaneous alternation test used to examine the Working memory. Spontaneous Alternation was calculated as the percentage of real alternations versus total alternation opportunities. ***C right,*** No differences were observed between control and IGF-IR^-/-^ mice in the Y-Maze Spontaneous Alternation Task, indicating a similar working memory (control=11, IGF-IR^-/-^ = 11, p=0,7474; Mann Whitney U Test).

